# Reconstruction of visual images from mouse retinal ganglion cell spiking activity using convolutional neural networks

**DOI:** 10.1101/2022.06.10.482188

**Authors:** Tyler Benster, Darwin Babino, John Thickstun, Matthew Hunt, Xiyang Liu, Zaid Harchaoui, Sewoong Oh, Russell N. Van Gelder

## Abstract

All visual information in mammals is encoded in the aggregate pattern of retinal ganglion cell (RGC) firing. How this information is decoded to yield percepts remains incompletely understood. We have trained convolutional neural networks with multielectrode array-recorded murine RGC responses to projected images. The trained model accurately reconstructed novel facial images solely from RGC firing data. In this model, subpopulations of cells with faster firing rates are largely sufficient for accurate reconstruction, and ON- and OFF-cells contribute complementary and overlapping information to image reconstruction. Information content for reconstruction correlates with overall firing rate, and locality of information contributing to reconstruction varies substantially across the image and retina. This model demonstrates that artificial neural networks are capable of learning multicellular sensory neural encoding, and provides a viable model for understanding visual information encoding.

**Significance Statement:** Convolutional neural networks can be trained on high-density neuronal firing data from the optic nerve to reconstruct complicated images within a defined image space.

## Introduction

Convolutional neural networks (CNN) achieve accurate performance across a wide variety of tasks in machine vision, including image recognition (*1)*, face generation (*2*), and instance segmentation (*3*). Although the original formulation of a CNN was inspired by the discovery of simple and complex cells in visual cortex (*4*), limitations on multiplex cell recording have hindered application of these models in reconstructing images from action potentials of the optic nerve. Early work on decoding visual stimuli from recordings of 20-30 RGC in the tiger salamander found no difference in performance between linear models and artificial neural networks (*5*). A recent study decoding an average of ∼80 RGC per reconstruction found that nonlinear models gave only minor improvements over a linear model (*6*), although, using simulated neural responses, it was suggested that a CNN would provide significant improvement over a linear model when recording from the entire retina (*7*).

Here, we utilize a 4096-channel microelectrode array (MEA) to record from thousands of spike-sorted units in an *ex vivo* preparation of the mouse retina. CMOS technology permits simultaneous 17kHz recording at a 42 µm electrode pitch and enables dense sampling of the mouse retina. A digital projector run through a high-resolution optical minimization path illuminates the retina at 1280×800 pixels, enabling the presentation of high-acuity stimuli. The inferior half of the retina is mounted on the MEA for each experiment. Stimuli are presented for 0.5 seconds, and spike sorted action potentials binned per unit per 100 ms for 1 second following stimulus onset (0.5 s of image and 0.5 s of dark recovery). These data are subsequently utilized by machine learning algorithms for decoding, in which the originally projected image is reconstructed using solely RGC spike data recorded by MEA (Figure 1A).

**Figure 1.**
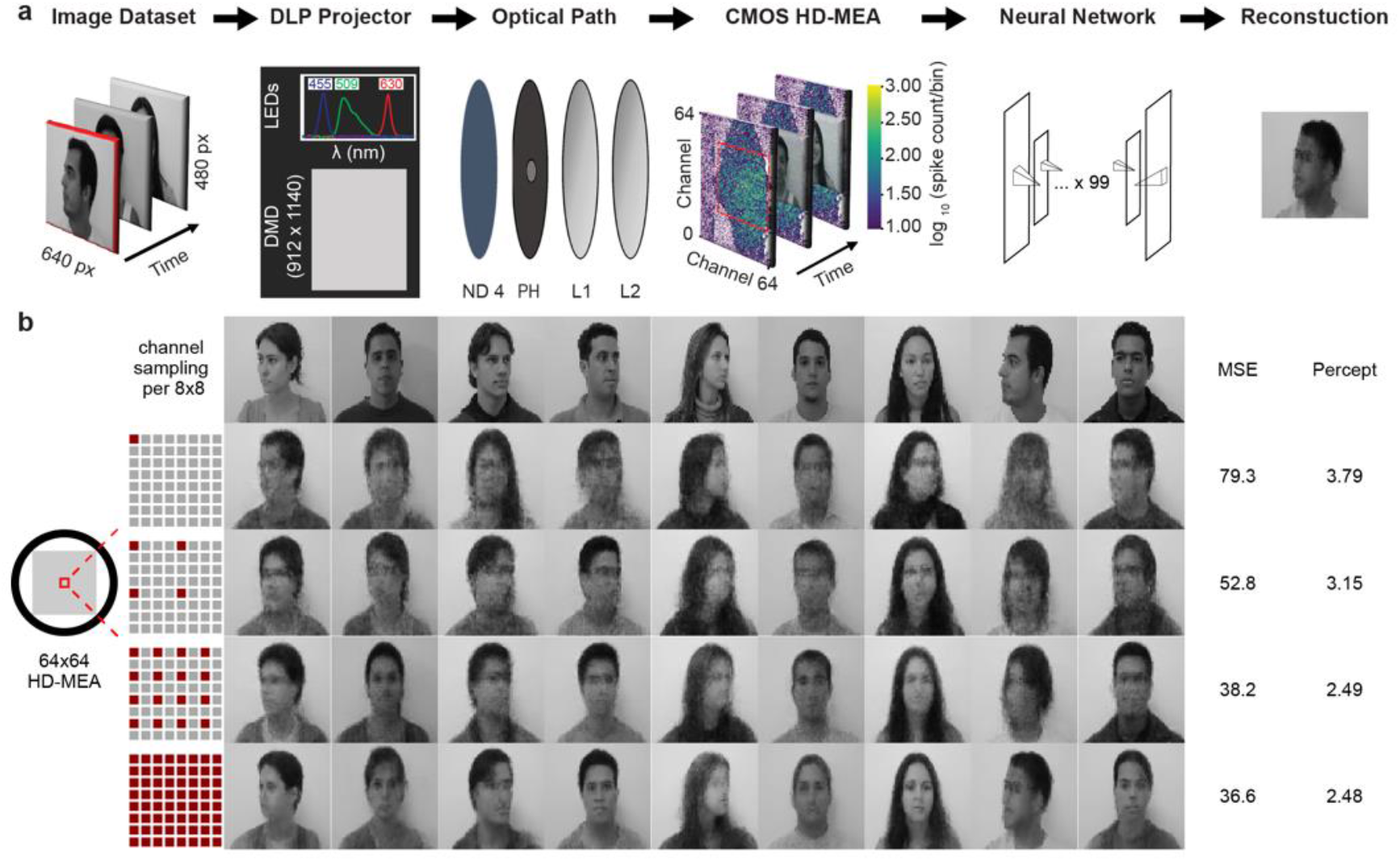
Neural network decoding of retinal ganglion cell spiking activity. **a**, Schematic overview of experimental method. A publicly available dataset of 640 × 480 pixel human facial images (FEI dataset, https://fei.edu.br/~cet/facedatabase.html) is projected onto an *ex vivo* retinal explant mounted, RGC-side down, on 4096 channel HD-MEA. Spike counts per 100ms for each spike-sorted unit are input to a U-net style residual network with 101 convolutional layers. The network is trained on 80% training set of 1400 images, and then used to predict the previously unused 20% of images (10% validation and 10% held-out test) using the RGC action potential train resulting from each image. **b**, Reconstruction accuracy improves with spatial resolution of electrode sampling. To evaluate reconstruction performance as a function of spatial sampling, models were trained on subsets of the recorded electrode channels, sampling every eighth electrode (8×8), every fourth electrode (16×16), every other electrode (32×32), or all electrodes (64×64). Top row of images is original projected image, lower rows are neural net reconstructions based on RGC MEA firing patterns. (MSE, mean square error; Percept, perceptual loss function; DLP, digital light processing; DMD, digital micromirror device; ND, neutral density filter; CMOS HD-MEA, complementary metal-oxide-semiconductor high-density multielectrode array; px, pixel; L1 and L2, condensing lenses 1 and 2 respectively; PH, pinhole).

## Results

### Image reconstruction varies perceptually by model architecture

Viability of murine retinal tissue *ex vivo* limits recording sessions to ∼6 hours, necessitating use of a limited image training set. As facial recognition has been extensively studied in machine vision (*8*), we selected a monochromatic version of the Fundação Educacional Inaciana (FEI) Face Database (*9*) of 200 individuals with 14 photos per person, including profile rotations up to 180 degrees, for a total of 2800 images. 2240 of these poses were randomly selected and presented to the retina singly for the training set, and 280 images each were held out and utilized as validation and final performance test set. We parameterized the map between recorded retinal signals and visual stimuli with a simplified realization of the residual network, consisting of 101 2-layer convolutional residual blocks (*10*) operating on an 8×8 intermediate spatial representation. We downsampled the network inputs to 8×8 with a single-strided convolution, and upsampled the network outputs to 64×64 with a transposed convolution. The structure of the resulting network (hereafter referred to as ResUNet) can be compared to the U-Net (*11*), which also maps spatial inputs to spatial outputs, using a lower-dimensional spatial bottleneck in the intermediate representation. In contrast to the U-Net, we use a single pair of downsampling and upsampling operations, and no long-distance skip connections between the inputs and outputs. Training was performed to optimize overall pixel-level mean squared error (MSE) in the reconstructions. As shown in Figure 1B, the trained ResUNet was able to generate accurate images from MEA-derived retinal ganglion cell firing data for images not in the training set. The model was generally able to reconstruct key image features including head position, hairline, clothing brightness, and overall background brightness. The number of recording channels used for training and test set had a strong effect on reconstruction quality (Fig. 1B). Downsampling of MEA data resulted in gradual degradation of image reconstruction, while using all 4096 channels of the multielectrode array resulted in the highest quantitative and qualitative performance.

### Decoding performance is model-dependent and consistent with rate coding hypothesis

Results from ResUNet were compared to log-linear regression models and multi-layer perceptron (MLP) models (*12, 13*), as well as a wide 18-layer ResNet with 2-layer fully-connected residual blocks (ResMLP) (Fig 2). All models produced images with qualitative features of the original image, and all produced MSE error within a 10-point range of 33-43. However, the qualitative results of different model architectures varied substantially, with ResUNet capturing key features of the original image such as gaze direction and facial features. Using a perceptual loss measure (*14*) (which sums squared errors between pixels) as an error measure, the ResUNet network produced the most accurate images. In contrast to previous studies (*5, 6*), we found that nonlinear reconstruction outperformed linear reconstruction. We attribute this difference primarily to choice of model architecture, as here we utilized standard approaches from machine learning methodology as opposed to adding a scalar nonlinear transform analogous to the linear-nonlinear (LN) encoding model. Our choice of dataset may also contribute, since constraining images to a known set of individuals may enable nonlinear models to impute missing data based on an internal classification of pose and person.

**Figure 2.**
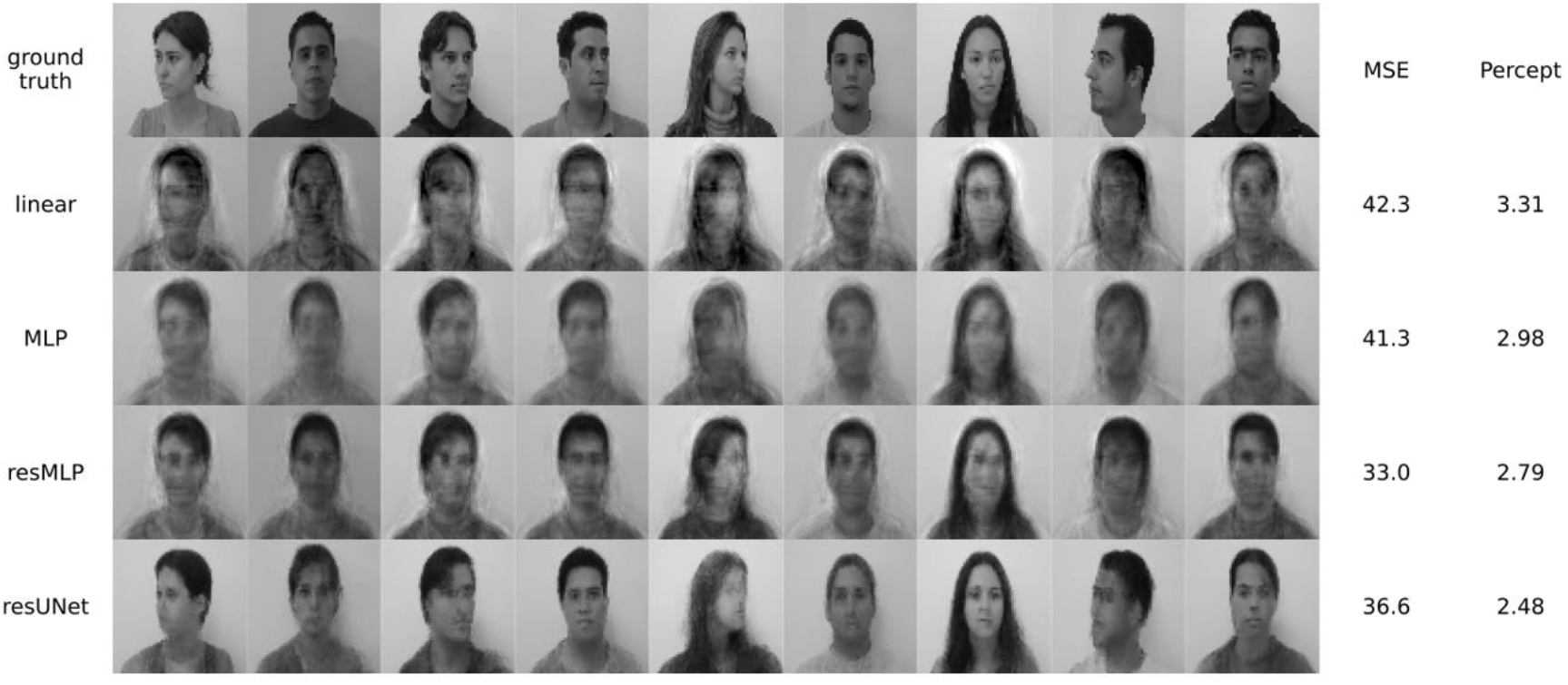
Neural net performance varies with model architecture. Representative original images from the FEI dataset (top row) and predicted images from four models: a linear model, a multilayer perceptron (MLP), an MLP using the residual block architecture (resMLP), and a U-net style architecture with residual network blocks of convolutions (resUNet). MSE, mean square error; Percept, perceptual loss function (see text).

The initial model utilized the full 1 s of image information per example (consisting of 0.5 s presentation and 0.5 s of dark recovery), divided into 100 ms bins. To determine the temporal distribution of information for reconstruction utilized by ResUNet, we first compared MSE for the model using solely the first or last 0.5 s. MSE for the first half second of presentation (32.9, green line in Figure 3) was nearly equivalent to that derived from the full 1 s presentation (31.1, green line in Figure 3), while the second 0.5 s resulted in a higher MSE (41.5, purple line) that was still superior to baseline (untrained) error (111.5, red line). The ability to generate an image during the dark half of image presentation is principally due to activity of OFF-cells (see below). To further determine the temporal distribution of information, each 1-second training image presentation was divided into 20 nonoverlapping 50-ms time-bin epochs. 20 separate models were trained, each using data from a single epoch. Firing data from each time bin epoch was represented as the sum of all spikes in that time period per retinal ganglion cell. MSE varied with time after image presentation, with two local minima in error, occurring at approximately 100-150 ms after image-on and image-off, respectively. This corresponded to the two peaks of firing activity with image on and off. However, the error from each of these time bins was greater than that found using the entire 1 s or first 0.5 s presentation, suggesting that most, but not all, information necessary for optimal reconstruction resides within these specific 50 ms bins or maximal firing activity.

**Figure 3.**
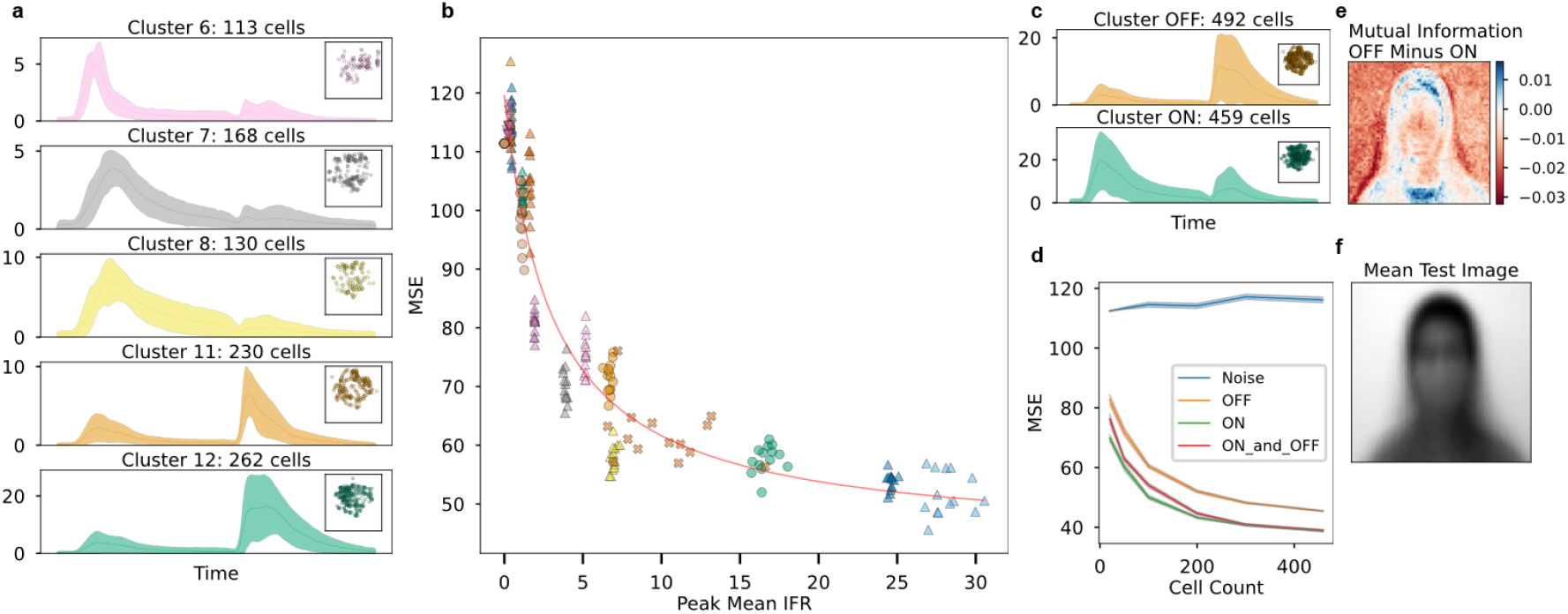
Contribution of RGC subtypes to image prediction accuracy. **a**, Mean instantaneous-firing-rate (IFR) +/- standard deviation for 5 example RGC clusters based on UMAP clustering. Inset shows spatial locations of units on MEA. **b**, MSE performance of 101 cells randomly selected from of each of 13 clusters, plotted as a function of to the maximum mean IFR per cluster. Fit with inverse proportional model. **c**, Firing pattern of grouped clusters assigned to ON-vs-OFF archetype. **d**, MSE performance +/- SEM of equal sampling of ON, OFF, ON+OFF, and noise cells. **e**, Difference in mutual information between ground truth and reconstruction for OFF versus ON cells (using 459 cells of each) across all test images. Positive values (blue) indicate regions where OFF cells have higher mutual information (and thus lower MSE). **f**, Mean of all held-out test projected images for reference for (e).

The murine retina has a diverse set of retinal ganglion cell types, characterized by their morphologic and immunohistochemical characteristics as well as firing responses to light stimuli (*15-18*). The contributions of different cell types to visual processing remains incompletely understood. To determine the relative contributions of different RGC types to reconstruction, we grouped RGCs based on their responses to image stimuli, followed by image-reconstruction performance by individual clustered cell type. Prior to clustering, the raw MEA signals were spike-sorted and smoothed to obtain the instantaneous firing rate (IFR) over time for each RGC. Averaging the IFR over all image presentations gave a feature vector of mean IFR at each time-point per RGC. Uniform manifold approximation and projection (UMAP) (*19*) was used to reduce the dimensionality of these feature vectors, and OPTICS unsupervised clustering (*20*) was performed to identify groups of cells. ‘Noise’ RGCs without a clear response to stimuli were identified by initial clustering. Following redaction of these cells, remaining cells were re-clustered to obtain the final groups. While most clusters showed some transient response to both stimulus presentation and cessation, the mean IFR tracings (across all cells and all image presentations) of these final clusters could be manually assigned to “ON” or “OFF” cells based on the inspection of tracings for predominant response. Figure 4A shows the average mean IFR +/- SEM for five selected cell types, along with their corresponding locations on the MEA array. These cells were generally homogenously distributed across the retina (while the majority of the “Noise” cells were distributed outside of the retina’s location on the MEA). Cells located near the center of the retina tended to have higher firing rate than cells distributed more peripherally. The performance of each cluster on facial-image reconstruction in terms of MSE is shown in Figure 4B. Because the smallest cluster had 101 cells, each point represents a single model trained on 101 randomly sampled RGCs from a single cluster. MSE loss was found to be inversely proportional to peak mean IFR across all cell types.

**Figure 4.**
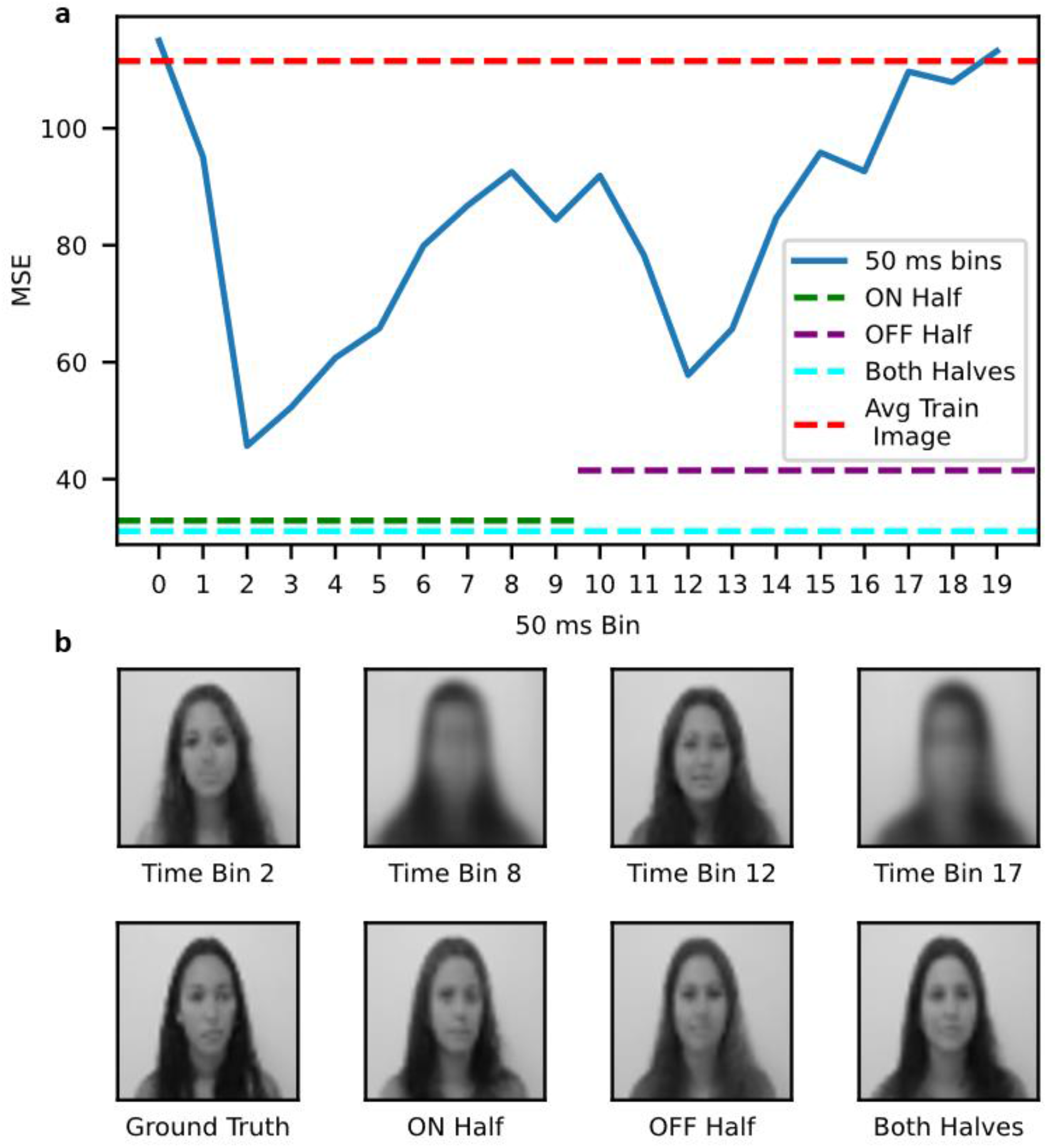
Neural net reconstruction accuracy varies with time after image exposure. **a**. Test-set MSE performance for models trained using firing data from each of 20 single 50-ms time bins. Performance for models trained on the entire first half (image) and second half (dark) of image presentations in green and purple, respectively. Teal represents model for total spike count for entire 1s epoch. The red line represents the MSE if the model always predicted the mean training image. **b**, Representative test-set reconstructions from FEI dataset are shown for models trained using 50 ms time bin 2, 8, 12, and 17 (top) and first half (ON, image presentation) and second half (OFF, image dark). Mean Train Image = average of all training images.

Clusters 8, 9, and 10 were selected as high-firing-rate “ON” cells, and clusters 11 and 12 were selected as high-firing-rate “OFF” cells. The aggregate mean IFR of the ON and OFF cells is shown in Figure 4C. Multiple models were trained using only either ON or OFF cells and with access to either the firing data from the first half of image exposure (face image shown) or the second half of image exposure (darkness between presentations). Each cell type performed best when given its respective image-exposure half. Models trained using only first-half ON-cell data and second-half OFF-cell data obtained mean MSE scores ± SEM of 41.1 ± 0.2 and 52.2 ± 0.3, respectively. In contrast, the cell type and exposure-half mismatched models trained using only second-half ON-cell data and first-half OFF-cell data obtained mean MSE scores ± SEM of 71.7 ± 0.5 and 63.7 ± 0.4, respectively. The performance of the ON and OFF cells was evaluated as the number of training RGCs randomly sampled from each increased, with Noise cells serving as a control. Noise cell performance did not improve as the number of cells increased (Figure 4D). Images reconstructed exclusively from ON cells had lower MSE than images constructed from solely OFF cells, and an even sampling of ON and OFF cells performed similarly to ON cells alone (Figure 4D). The per-pixel mutual information (MI) between predicted and ground truth pixels was calculated across all images in our test set for both ON-cell-only models and OFF-cell-only models and mapped on the mean test set image (Figure 4E). MI performance varied by cell type across the image, with ON cells encoding more information about the background, while OFF cells encoded more information for high-contrast regions like the outline of the head. This analysis suggests that, for the ResUNet model, ON- and OFF-cells encode partially redundant information for static images, but that reconstruction of specific features within that image may utilize more information from one or the other cell type depending on local image characteristics.

### Reconstructions respect retinotopy

We next sought to understand how local cell spiking information is used in reconstructions. We used saliency mapping (*21*) to answer the following question: what parts of the recorded retina signals contribute to different patches of the image, such as an eye in a reconstructed face? We computed the gradient of reconstructed output (image) with respect to input (RGC firing); the intensity of this saliency map represents the output change with respect to a small perturbation in the input. We calculated the gradient of two particular patches from a single test-set image (Figure 5A) with respect to the recorded retina signal and plotted the absolute value of the gradient for each of the 4096 channels, averaging over the spike-sorted units, using a reconstructed image from the training set. Figure 5B shows a saliency map averaged over all data in the test-set, keeping the location of the patches identical as in Figure 5A. A similar analysis performed on a per-pixel basis is shown in Supplementary Video 1. Local spiking information is used for local image reconstruction, although the area utilized is substantially larger than receptive field size of single retinal ganglion cells. To determine overall locality of information contribution to reconstruction, we measured how much of the saliency score is captured in a local neighborhood of the patch (Figure 5C). Increasing the area of a circle centered at the middle of the patch (normalized by 4096, the total area of the input signal) and measuring the resulting local saliency score summed over all the retinal signal positions inside of that circle (normalized by the total saliency score summed over all 4096 input retinal signal positions) demonstrated that 50% of the saliency score is captured by 28.2% of the input positions. We also examined the locality of the influence each retinal signal has on the reconstructed image. Figure 5D shows the saliency of the reconstructed faces for each position (channel) in the recorded retinal signal. We selected several small patches in the output image and determined how the input retinal signal contributes to the particular patch. Supplementary Videos 1 and 2 demonstrate an equivalent analysis done on a per-channel basis. Again, local information is used preferentially in reconstruction, but regions contributing to the reconstruction may be found a substantial distance from the reconstructed patch. In aggregate, the saliency analysis demonstrates that the neural network is combining local retinal firing information with an imputed expectation of typical features common to the set of training images to produce the final image. A similar process is thought to underlie human visual cognition (*22*).

**Figure 5.**
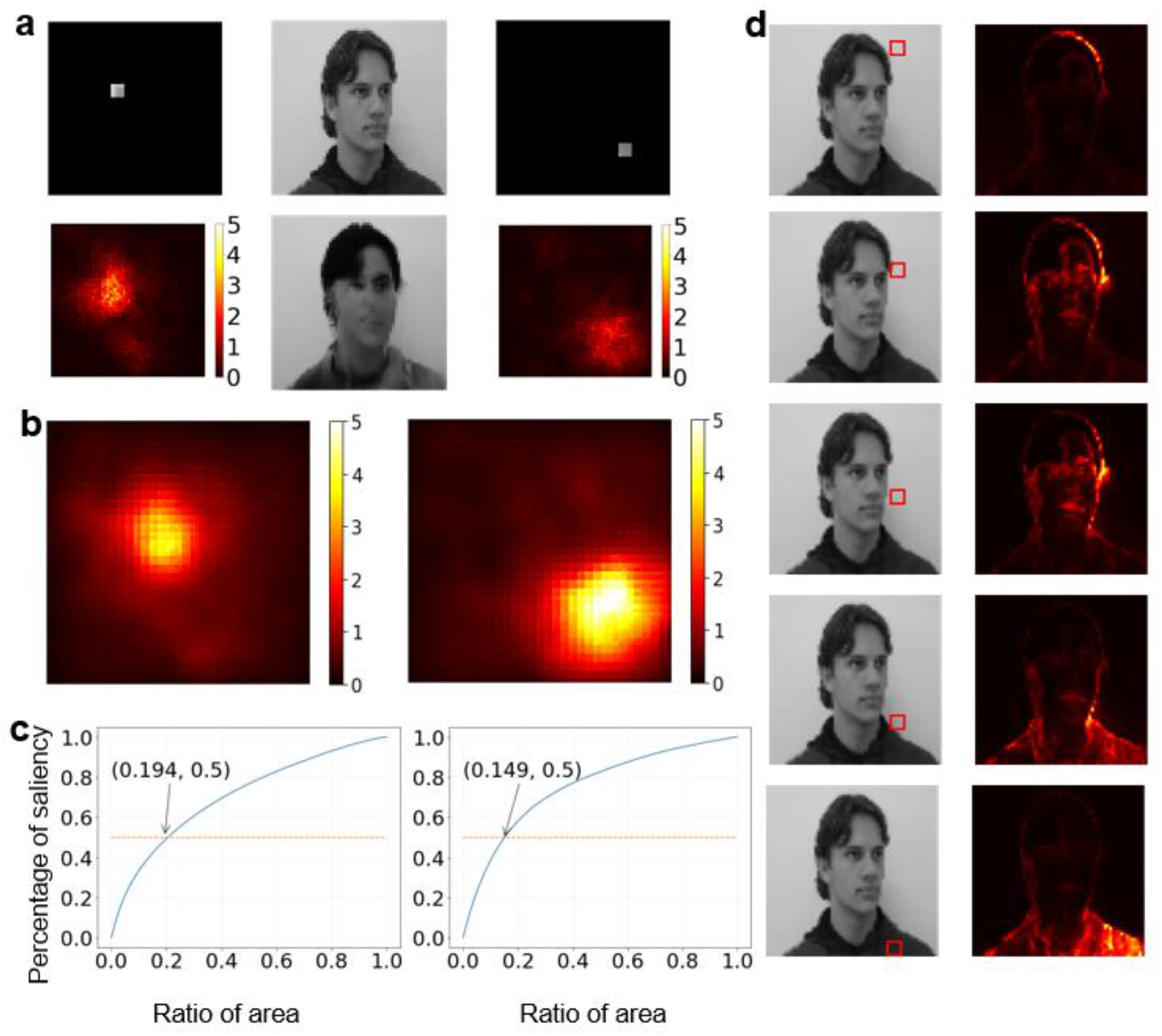
Saliency map for neural net image reconstruction. **a**. Representative example from DEI dataset of MEA channels contributing to local reconstruction. Top shows two image patches of interest (squares) on ground truth image (center). Heat map reflects relative contribution of MEA locations to reconstruction (center). **b**. Mean saliency map across all test images. **c**. Saliency as a function of locality. A ‘ball’ of image space is located at each point of interest in (a), and saliency measured with expansion of this ball, shown as fraction of total image area (blue). 50% line demonstrates ball area at which half of saliency is achieved. **d**. Reverse saliency analysis with heat map demonstrating all reconstructed pixels affected by specific MEA location (red box).

Finally, we examined the extent to which continuous local information is utilized by the model for image reconstruction. ResUNet was again trained on the facial image dataset, but this time, either the image was scrambled by lossless perturbation, the locations of RGC firing on the array was scrambled by the same algorithm, or both (Supplementary Figure 1). We first applied ResUNet to the task of unscrambling the original image. The net was able to perform this task with MSE of ∼10. We next trained ResUNet using scrambled images as output, and then unscrambled the resulting images by reversing the lossless scrambling algorithm. Relative to baseline MSE (38.2), the resulting reconstructions had much higher MSE (72.6). The same result occurred when ganglion cell position on the MEA was scrambled, with MSE of 75.2 Scrambling both image and RGC array position resulted in worse MSE than either perturbation alone (MSE 88.2). Taken together with the previous saliency analysis, this demonstrates that ResUNet makes use of continuity and local correlation of image and RGC firing in generating reconstructions.

## Discussion

Here we show that a convolutional neural network can be trained on high-density neuronal firing data from the optic nerve to reconstruct complicated images within a defined image space. We find that information used for reconstruction is correlated with instantaneous firing rate; that different retinal ganglion cell subtypes contribute different information to the reconstruction in this model, generally in proportion to peak firing rate; and that information is preferentially (but not exclusively) used locally, with larger areas of influence than would be predicted by classical cellular receptive fields. Machine learning approaches have been successfully employed in numerous image-transformation domains over the past several years (*2*) but to our knowledge this is the first successful use of machine learning to convert biologically encoded neural firing information from a sensory nerve into image generation, which is the essence of vision. The resulting model results in more accurate reconstructions of from data than are obtained from linear models (*23*).

Surprisingly, we find that while the ResMLP model performs best in terms of MSE, the convolutional ResUNet architecture performs better in terms of perceptual loss. Qualitatively, the two models display different biases under uncertainty. While the ResMLP model tends to blur uncertain regions of an image, ResUNet tends to impute textures that were seen during training. When information is severely limited, this can lead to perceptually plausible but jarring mistakes, as in the lowest sampling condition for Figure 1, where several reconstructions mistakenly depict a person with long hair whereas the original image displays short hair. However, when utilizing the full array, the reconstructions of the ResUNet appear remarkable even compared to ResMLP, as in Figure 2, reconstruction 7, where facial features like the eyes, eyebrows, nose, and mouth are sharply defined and well-matched to the displayed image. Although regurgitation of training data is a hallmark of overfitting, the ResUNet model nonetheless generalizes better than the other models in terms of perceptual loss in withheld test images. Higher order cortical processing underlies human facial recognition, and false recognition of unfamiliar faces or inability recognize familiar faces (prosopagnosia) are well-recognized sequalae of neurologic damage, particularly to the right occipitotemporal structures of the brain (*24*). It is possible that deep CNNs and human cortex employ similar processing to impute visual features from previously viewed faces under inconclusive sensory information.

At least part of the improvement provided by the CNN-derived model compared with linear models is due to its ability to impute expected features of human faces. The dataset used for training was intentionally limited given the relatively short period of training time available on the *ex vivo* mouse retina (∼6 hours). It is apparent the CNN derives an average image (or ‘Eigenface’) which will be produced when limited data are available. However, saliency analysis demonstrates that local information is used extensively in reconstruction within this space. In this way, the CNN mirrors processes thought to underlie visual percept generation (*4*), in which expected image may drive percept from equivalent visual data (as manifest, for example, in the well-studied Rubin’s goblet ‘vase vs. faces’ optical illusion (*25*)).

In analyzing the relative contributions of retinal cell types to reconstruction, we confirm that ON-cells contribute primarily to reconstruction during the initial image presentation, and OFF-cells utilize information primarily from the period immediately after image cessation. While the information provided by these two channels are partially redundant (and good quality reconstruction can be generated by either alone), mutual information analysis suggests that different portions of the static image rely preferentially on one channel or another, with background brightness, for instance, dependent primarily on ON-channel function. We also find that individual RGC types appear to be sufficient for reconstruction of images, consistent with the parallel information model (*26*). Further, in analyzing RGC subtypes, we find that the contribution of individual subtypes to image reconstruction appears to vary with peak firing rate, suggesting that rate coding (as opposed to temporal pattern encoding (*27*)) may be sufficient for high fidelity information transfer to the brain in the context of static images. Moreover, we found that a model trained on single 1 s time bin per spike-sorted unit led to better generalization than a model trained with 10 × 100 ms time bins per spike-sorted unit, further supporting the rate coding hypothesis.

Absence of a testable model for ensemble integration of optic nerve firing into high resolution images has precluded testing of important questions in retinal and visual physiology. The ResUNet model allows for assessment of the necessity and sufficiency of specific cell and cell-class information in image reconstruction within the model. We find that both ON- and OFF-cells contribute partially redundant information to image reconstruction, but that neither is strictly necessary for good quality static image reconstruction. This result is consistent with results seen in individuals lacking mGluR6 function, which eliminates ON-bipolar function and ON-cell activation. Such subjects have poor visual responses under dim light but show near-normal central visual acuity in photopic conditions despite loss of the ON-pathway (*28*), as is seen in the OFF-only results in the ResUNet model.

The present work represents a first application of CNN methods to decoding of complex sensory nerve signaling. The test case of monochromatic facial images is limited, and many extensions will be possible, including incorporation of a wider variety of images from different natural scenes; incorporation of color; and incorporation of motion. Further, the current studies were done utilizing training and validation on single retinas. The extent to which ensemble networks trained on multiple retinas simultaneously can be used to decode the output from novel retinal signals remains to be determined. Finally, the current work was limited to murine retina. MEA recording is feasible in non-human primate and human retina (*29*) and may allow for decoding of human vision from retinal activity.

Several technologies, including bio-electronic retinal chip prostheses (*30*), opsin-based inner retinal gene therapy (*31*), and small molecule photoswitches (*32*) have the potential to confer light-dependent firing activity on retinal ganglion cells in individuals blind from outer retinal degeneration caused by macular degeneration or hereditary retinal degeneration (*33*). The encoding of the visual world with these prosthetics will not be native. Development of CNN-based models for visual reconstruction from optic nerve will allow for prediction of percepts from prosthetic encoding and, more importantly, may allow for modification of prosthetic encoding to more faithfully recreate natural-appearing percepts.

## Materials and Methods

### Animals

C57BL/6J (*wt*) mice between 6-12 months old (Jackson laboratory, Bar Harbor, ME, USA) were used in this study. Both male and female mice were incorporated into studies randomly. All animals were treated in accordance with the ARVO Statement for the Use of Animals in Ophthalmic and Vision Research and all word done under approved IACUC protocol at the University of Washington.

### Stimulator

A DLP LightCrafter 4500 projector (Texas Instruments, Dallas, TX, USA) coupled to a custom optical, two-lens system capable of 6 cpd resolution focused light stimulation onto the retina from below (i.e. presenting to ganglion cell side of retina, consistent with normal vision). The system provided 1280×800 pixels of spatiotemporally patterned stimulation over the area of the MEA with a refresh rate of 60 Hz and control of brightness and simultaneous RGB LED operation. The DLP projector is outfitted with 3 independently controlled RGB LEDs with a recorded maximum emission at 630, 509 and 455 nm for red, green and blue respectively with an average half-width of ± 30 nm. For white light stimulations, equivalent photon flux (∼ 3.5e+13 photons/cm^2^/s) per LED was used.

### Dataset

Projected images were taken from the FEI Face Database (https://fei.edu.br/~cet/facedatabase.html), a publicly available database of standardized facial images. The dataset includes 14 images for each of 200 individuals (100 male and 100 female), for a total of 2800 images. All images are taken against a white homogenous background in an upright frontal position with profile rotation of up to about 180 degrees. Each image is 640×480 pixels. All participants gave informed consent for use of their images in the public dataset, and images are not copyrighted.

### MEA Recordings

Euthanasia of all mice was performed using CO_2_ narcosis followed by cervical dislocation. Eyes were rapidly enucleated and placed into room-temperature artificial cerebral spinal fluid (ACSF) under dim red light. ACSF solution contained: 125 mM NaCl, 2.5 mM KCl, 1.25 mM NaH_2_PO_4_, 1 mM MgCl_2_, 2 mM CaCl_2_, 26 mM NaHCO_3_, and 20 mM D-glucose and aerated with 95% O_2_/5% CO_2_. Isolated retinas hemi-sectioned into superior and inferior retina, were placed whole-mount with retinal ganglion cell layer facing down, onto a 4096-channel HD-MEA Arena chip and recorded on BioCAM X system (3Brain, Wadenswil, Switzerland). The electrode recording area spans 2.67 mm x 2.67 mm. Retinas were kept at 34°C and continuously perfused with ACSF at 3 mL/min and allowed to settle and recover for at least 1 h prior to recordings. Detection and processing of extracellular spikes and waveforms was performed using BrainWave 5 (3Brain, Wadenswil, Switzerland). Spikes were high-pass filtered at 300 Hz and digitized at 18 kHz with a spike threshold setting of 5 SD for each channel. Single-unit spikes were sorted using custom software using Herdingspikes2 (*34*).

## Data / Code availability

Data and code used for analysis in this paper is available at https://github.com/tbenst/2021-retina-reconstructions.

## Funding

National Institutes of Health grant K99EY03133301 (DB), P30EY001730 (RVG), an unrestricted grant from Research to Prevent Blindness, and the Mark J. Daily Research Fund.

## Notes

### Competing Interest Statement

The authors have declared no competing interest.

## References

1. K. He, X. Zhang, S. Ren, J. Sun. Identity Mappings in Deep Residual Networks. In: Leibe, B., Matas, J., Sebe, N., Welling, M. (eds) Computer Vision – ECCV 2016. ECCV 2016. Lecture Notes in Computer Science(), vol 9908. Springer, Cham.

2. T. Karras, S. Laine, T. Aila, A Style-Based Generator Architecture for Generative Adversarial Networks (2018)

3. K. He, G. Gkioxari, P. Dollár, R. Girshick. Mask R-CNN. In Proceedings of the IEEE international conference on computer vision, 2961–2969 (2017).

4. K. Fukushima, Neocognitron: A self-organizing neural network model for a mechanism of pattern recognition unaffected by shift in position. Biological Cybernetics 36, 193–202 (1980).

5. D. K. Warland, P. Reinagel, M. Meister, Decoding visual information from a population of retinal ganglion cells. Journal of neurophysiology 78, 2336–2350 (1997).

6. N. Brackbill et al., Reconstruction of natural images from responses of primate retinal ganglion cells. eLife 9, 1–65 (2020).

7. N. Parthasarathy, E. Batty, W. Falcon, T. Rutten, M. Rajpal, E.J. Chichilnisky, L. Paninski. Neural networks for efficient bayesian decoding of natural images from retinal neurons. Advances in Neural Information Processing Systems, 30 (2017).

8. F. Schroff, D. Kalenichenko, J. Philbin, J. FaceNet: A unified embedding for face recognition and clustering. IEEE Conference on Computer Vision and Pattern Recognition (CVPR),815–823 (2015).

9. C. E. Thomaz, G. A. Giraldi, A new ranking method for principal components analysis and its application to face image analysis. Image and Vision Computing 28, 902–913 (2010).

10. K. He, X. Zhang, S. Ren, J. Sun, Deep Residual Learning for Image Recognition. arXiv e-prints, arXiv:1512.03385 (2015).

11. O. Ronneberger, P. Fischer, T. Brox, U-Net: Convolutional Networks for Biomedical Image Segmentation. 2015.

12. F. Rosenblatt, The perceptron: a probabilistic model for information storage and organization in the brain. Psychol Rev 65, 386–408 (1958).

13. D. E. Rumelhart, G. E. Hinton, R. J. Williams, Learning representations by back-propagating errors. Nature 323, 533–536 (1986).

14. J. Johnson, A. Alahi, L. Fei-Fei, Perceptual Losses for Real-Time Style Transfer and Super-Resolution. arXiv e-prints, arXiv:1603.08155 (2016).

15. T. Baden et al., The functional diversity of retinal ganglion cells in the mouse. Nature 529, 345–350 (2016).

16. K. Farrow, R. H. Masland, Physiological clustering of visual channels in the mouse retina. J Neurophysiol. Apr;105(4):1516–30 (2011)

17. J. Jouty, G. Hilgen, E. Sernagor, M. H. Hennig, Non-parametric Physiological Classification of Retinal Ganglion Cells in the Mouse Retina. Frontiers in Cellular Neuroscience 12, (2018).

18. Y. Zhang, I.-J. Kim, J. R. Sanes, M. Meister, The most numerous ganglion cell type of the mouse retina is a selective feature detector. Proceedings of the National Academy of Sciences 109, E2391–E2398 (2012).

19. L. McInnes, J. Healy, J. Melville, UMAP: Uniform Manifold Approximation and Projection for Dimension Reduction. 2018.

20. M. Ankerst, M. M. Breunig, H. P. Kriegel, J. Sander, OPTICS: ordering points to identify the clustering structure. SIGMOD Rec. 28, 49–60 (1999).

21. K. Simonyan, A. Vedaldi, A. Zisserman, Deep Inside Convolutional Networks: Visualising Image Classification Models and Saliency Maps. 2013.

22. C. Summerfield, T. Egner, Expectation (and attention) in visual cognition. Trends in Cognitive Sciences 13(9), 403–409 (2009).

23. S. Nirenberg, C. Pandarinath, Retinal prosthetic strategy with the capacity to restore normal vision. Proceedings of the National Academy of Sciences 109, 15012–15017 (2012).

24. J. J. Barton, Disorders of face perception and recognition. Neurol Clin. May;21(2):521–48 (2003)

25. M. Takashima, K. Fujii T Fau - Shiina, K. Shiina, Face or vase? Areal homogeneity effect.

26. J. Gjorgjieva, H. Sompolinsky, M. Meister, Benefits of Pathway Splitting in Sensory Coding. The Journal of Neuroscience 34, 12127–12144 (2014).

27. R. Gütig, T. Gollisch, H. Sompolinsky, M. Meister, Computing Complex Visual Features with Retinal Spike Times. PLoS ONE 8, e53063 (2013).

28. T. P. Dryja et al., Night blindness and abnormal cone electroretinogram ON responses in patients with mutations in the GRM6 gene encoding mGluR6. Proc Natl Acad Sci U S A 102, 4884–4889 (2005).

29. K. Reinhard, T. A. Münch, Visual properties of human retinal ganglion cells. PLOS ONE 16, e0246952 (2021).

30. J. O. Mills, A. Jalil, P. E. Stanga, Electronic retinal implants and artificial vision: journey and present. Eye (Lond). 2017 Oct;31(10):1383–1398.

31. M. Ratner, Light-activated genetic therapy to treat blindness enters clinic. Nature Biotechnology 39, 126–127 (2021).

32. R. N. Van Gelder, Photochemical approaches to vision restoration. Vision Research 111, 134–141 (2015).

33. B. Roska, J. A. Sahel, Restoring vision. Nature 557, 359–367 (2018).

34. J.-O. Muthmann et al., Spike Detection for Large Neural Populations Using High Density Multielectrode Arrays. Frontiers in Neuroinformatics 9, (2015).

