## Supplementary material for "Reconstruction of visual images from mouse retinal ganglion cell spiking activity using convolutional neural networks": Materials and methods, supplementary figure

**This PDF file includes:**

Fig. S1
Captions for Movies S1 to S2

**Other Supplementary Materials for this manuscript include the following:**

Movies S1 to S2

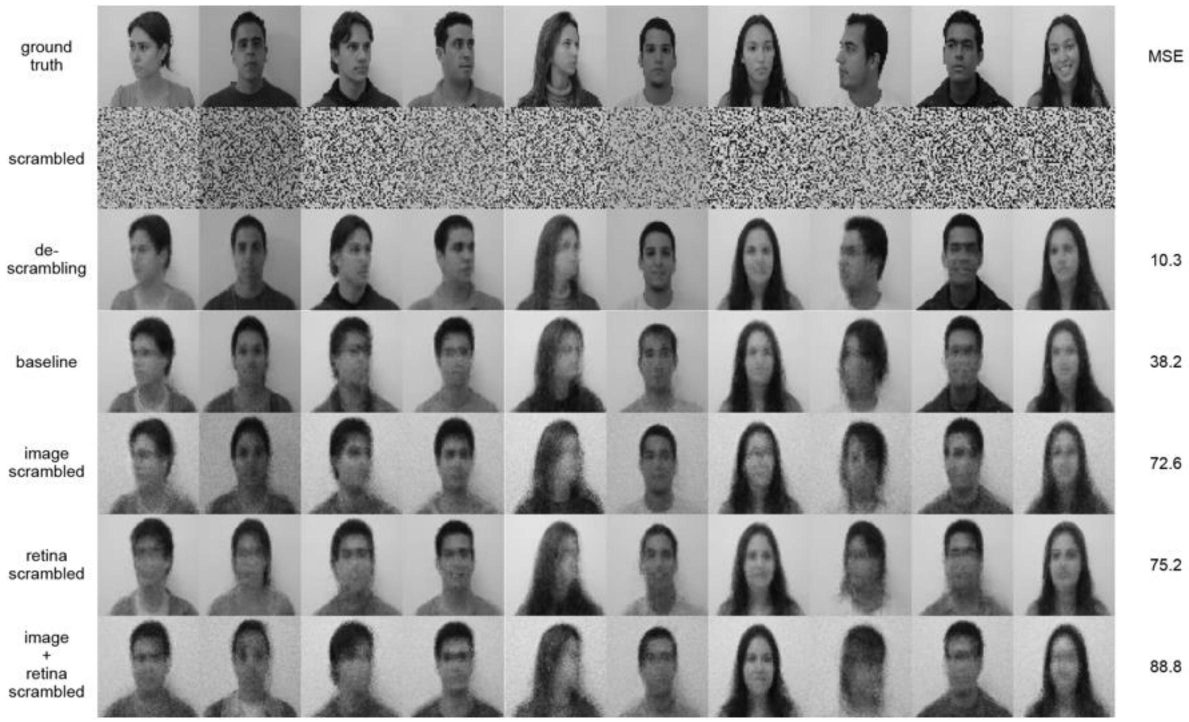


**Fig. S1.**

Row 1 ('ground truth'): a sample of FEI dataset images presented to the retina. Row 2 ('scrambled'): the same images, with the pixel locations scrambled according to a fixed random permutation. Row 3: the predictions of a ResUNet model trained to predict the original images (Row 1) given the scrambled images as input (Row 2). This baseline demonstrates the limits of the ResUNet to learn a permutation of spatial data, given losslessly permuted image as input. Row 4 ('baseline'): the predictions of a ResUNet model trained to predict original images, given retina data as input. Row 5 ('image scrambled'): the predictions of a ResUNet model trained to predict scrambled images, given retina data as input. This removes the spatial structure from the target images, and the performance of the ResUNet is substantially degraded. Row 6 ('retina scrambled'): the predictions of a ResUNet model trained to predict original images, given scrambled retina data as input. This removes the spatial structure from the retina, and the performance of the ResUNet is similarly degraded. Row 7('image + retina scrambled'): the predictions of a ResUNet model trained to predict scrambled images given scrambled retina data as input. Both inputs and outputs are scrambled according to the same spatial permutation, but this further degrades the performance of the ResUNet model.

**Movie S1.**

**Visualization of projective field from single pixel to MEA channels**. With the average image as background, the mouse points to a seed pixel, and colors indicate a positive influence or negative influence of spike count per electrode on the seed pixel reconstruction. To calculate these values, a one-hot gradient mask was backpropagated through a 32x32 channel ResUnet model. For an individual photo, this value represents the change in spike count per time bin per spike-sorted unit that would need to change for the pixel value to brighten for the seed pixel only. Gradients were averaged across all test images, and summed across time bins and spike-sorted units. This gives the linear approximation of the average projective field from each pixel to all MEA channels.

**Movie S2**

**Visualization of receptive field from single pixel to MEA channels**

With the average image as background, the mouse points to a seed MEA channel, and colors indicate a positive correlation or negative correlation of pixel values on the spike count of the seed MEA channel. To calculate these values, we use the gradient values calculated as in supplementary video 1, but instead index by MEA channel instead of by pixel. For an individual photo, this value represents a local prediction that a small perturbation of the image by this gradient will only change the spike count on this individual MEA channel. When averaged, this gives a linear approximation of the average receptive field for each MEA channel, which includes up to six spike-sorted units.
